# Chromosomal instability increases radiation sensitivity

**DOI:** 10.1101/2024.09.13.612942

**Authors:** Pippa F. Cosper, Maha Paracha, Kathryn M. Jones, Laura Hrycyniak, Les Henderson, Ava Bryan, Diego Eyzaguirre, Emily McCunn, Elizabeth Boulanger, Jun Wan, Kwangok P. Nickel, Vanessa Horner, Rong Hu, Paul M. Harari, Randall J. Kimple, Beth A. Weaver

## Abstract

Continuous chromosome missegregation over successive mitotic divisions, known as chromosomal instability (CIN), is common in cancer. Increasing CIN above a maximally tolerated threshold leads to cell death due to loss of essential chromosomes. Here, we show in two tissue contexts that otherwise isogenic cancer cells with higher levels of CIN are more sensitive to ionizing radiation, which itself induces CIN. CIN also sensitizes HPV-positive and HPV-negative head and neck cancer patient derived xenograft (PDX) tumors to radiation. Moreover, laryngeal cancers with higher CIN prior to treatment show improved response to radiation therapy. In addition, we reveal a novel mechanism of radiosensitization by docetaxel, a microtubule stabilizing drug commonly used in combination with radiation. Docetaxel causes cell death by inducing CIN due to abnormal multipolar spindles rather than causing mitotic arrest, as previously assumed. Docetaxel-induced CIN, rather than mitotic arrest, is responsible for the enhanced radiation sensitivity observed in vitro and in vivo, challenging the mechanistic dogma of the last 40 years. These results implicate CIN as a potential biomarker and inducer of radiation response, which could provide valuable cancer therapeutic opportunities.

**Statement of Significance:** Cancer cells and laryngeal tumors with higher chromosome missegregation rates are more sensitive to radiation therapy, supporting chromosomal instability as a promising biomarker of radiation response.

## Introduction

Aneuploidy, a state of unbalanced chromosome content that differs from a multiple of the haploid, is a hallmark of cancer that is present to varying extents in ∼85% of solid tumors (1,2). While aneuploidy can result from a single abnormal mitotic division, continuous chromosome missegregation produces chromosomal instability (CIN), an ongoing rate of missegregation events over the course of multiple cell divisions which is present in approximately 45% of solid tumors (3,4). There are many potential causes of chromosome missegregation including weakened signaling from the mitotic spindle assembly checkpoint (5,6), hyperstable attachments between microtubules of the mitotic spindle and the microtubule attachment sites (kinetochores) on sister chromatids (7), defects in sister chromatid cohesion (8), centrosome amplification (9,10), replication stress (11), and ionizing radiation (12), among others. Whole chromosome missegregation results in numerical CIN (13), while missegregation of chromosome arms or fragments lead to structural CIN, which includes chromosomal alterations such as translocations and rearrangements due to DNA damage or replication stress (11). Both types of CIN result in aneuploidy.

Ionizing radiation (IR) is used to treat over 50% of cancer patients and is a well-known cause of CIN (reviewed in (14)), though this has not been well characterized in head and neck cancer. The main mechanism of radiation cytotoxicity is formation of double-stranded DNA breaks which can yield acentric fragments, chromosomal translocations, or dicentric chromosomes if repaired erroneously (15). These aberrant chromosomes are commonly missegregated during mitosis. Acentric fragments, which are unable to attach to microtubules of the mitotic spindle because they lack kinetochores, fail to congress to the spindle equator during metaphase, resulting in misaligned chromosomes (12). As cells progress through later stages of mitosis, acentric chromosome fragments lag behind the segregating masses of DNA and are randomly segregated. When the two kinetochores on dicentric chromosomes attach to microtubules from opposite spindle poles, the dicentric chromosome is stretched between the segregating DNA masses, forming a chromatin bridge. These bridges can be maintained beyond mitosis into the next G1 (16), or they can break resulting in a chromosome breakage-fusion-bridge cycle yielding extensive genomic rearrangements (17). In addition to effects on DNA, radiation can also cause centrosome amplification leading to abnormal multipolar spindles (18,19). Multipolar spindles that are maintained throughout mitosis typically result in inviable daughter cells due to extensive chromatin loss (9,20–22). Focusing multipolar spindles into near-normal bipolar spindles increases daughter cell survival but, at least in cases of centrosome amplification, is associated with increased lagging chromosomes (9,10). Radiation also induces lagging chromosomes by increasing the stability of kinetochore-microtubule attachments, which impairs the normal process of correcting erroneous attachments between kinetochores and microtubules emanating from the inappropriate spindle pole (7,23). Missegregated chromosomes and chromosome fragments sometimes form micronuclei after mitosis (24,25). Chromatin in micronuclei often undergoes DNA damage in the subsequent cell cycle leading to extensive genomic rearrangements in a process known as chromothripsis (26). Thus, ionizing radiation results in both structural and numerical CIN and aneuploidy.

In general, CIN in cancer is associated with poor prognosis, acceleration of tumor evolution, altered treatment response, immune evasion and metastasis by increasing genomic heterogeneity (22,27–31). However, CIN is not a dichotomous variable and the rate of CIN is what appears to dictate cell fate. Low rates of CIN (1-4 chromosomes every few divisions) can be tumor promoting due to gain of oncogenes or loss of tumor suppressors, although most aneuploid cells are not transformed (32–34). In contrast, higher rates of CIN lead to cell death and tumor suppression due to loss of essential chromosomes (20,35–40). Thus CIN can promote or suppress tumors, or do neither, depending on the level of CIN and tissue context (reviewed in (4,41)). Combining two sources of tolerable rates of CIN increases the total CIN over a maximally tolerated threshold leading to cell death and tumor suppression (35–37,42). This phenomenon has been demonstrated in both cell culture and murine models of different cancer types. Importantly, high rates of CIN can cause tumor suppression even in the case of oncogene-driven tumors (43). In agreement with this, high rates of CIN, as measured by centromeric fluorescence in situ hybridization (FISH), correlate with improved prognosis and survival in two independent cohorts of breast cancer patients (44,45). These data support the hypothesis that cancer cells with higher levels of CIN at baseline are more sensitive to radiation since they are closer to their maximally tolerated threshold of chromosome loss. Indeed, rectal adenocarcinomas with higher levels of anaphase defects showed improved pathological response to neoadjuvant chemoradiation therapy (46). In accordance, suppression of CIN increased radiation resistance in a murine glioma model (12), likely by maintaining CIN well below the maximally tolerated threshold. Thus, there appears to be a tolerable range of CIN in cancer cells; increasing this above a maximally tolerated threshold leads to excessively high chromosome missegregation and cell death.

Currently, there is no clinically approved method to predict radiation response or resistance. Outside of clinical trials, it is common to treat cancers of the same type with the same radiation dose. This is despite the fact that head and neck cancers positive for Human Papillomavirus (HPV) have improved outcomes relative to their HPV-negative counterparts (47,48). Radiation to the head and neck often leads to long term side-effects that impair quality of life including dry mouth and difficulty swallowing and speaking. A reliable predictive biomarker could improve outcomes in these patients by allowing dose de-escalation in select patients, permitting successful cancer therapy with fewer complications. Furthermore, a predictive biomarker would identify patients likely to relapse or incompletely respond, who could benefit from escalated therapy, such as addition of a radiosensitizer.

Docetaxel has long been recognized as a radiosensitizer in preclinical models (49–52) and clinical trials (53–58), and is a component of definitive treatment in multiple cancer types, including head and neck cancer. Docetaxel is a semi-synthetic analogue of paclitaxel; both are chemotherapeutics that stabilize microtubules by promoting polymerization of tubulin subunits (59–61). It is well known that both docetaxel and paclitaxel cause mitotic arrest at high concentrations (49,62). Mitotic arrest was generally accepted as the anti-cancer mechanism of microtubule-targeting agents for decades. However, direct sampling of breast cancers 20 hours after paclitaxel treatment revealed substantially lower drug levels in tumors than traditionally used experimentally (20,22,63). Importantly, these low levels of paclitaxel did not cause mitotic arrest, but instead resulted in formation of multipolar spindles. Persistence of multipolar spindles throughout mitosis caused cell death due to lethal rates of CIN. Increasing CIN over the maximally tolerated threshold on multipolar spindles, rather than mitotic arrest, was shown to be the mechanism of breast cancer cell death induced by paclitaxel and other microtubule-targeted agents in cells and patient tumors (20–22,63). The presumed mechanism of radiosensitization by docetaxel has been induction of mitotic arrest (50–52), as mitosis is the most radiosensitive phase of the cell cycle. However, docetaxel may instead act similarly to paclitaxel by inducing CIN on multipolar spindles rather than mitotic arrest.

Here, we use engineered isogenic CIN and non-CIN cell line pairs, patient-derived xenografts, and patient biopsies to show that baseline CIN sensitizes head and neck and cervical cancer cells to radiation therapy. Additionally, we reveal a novel mechanism of radiosensitization by docetaxel: rather than inducing cell death as a consequence of mitotic arrest, docetaxel increases CIN due to multipolar spindles. Our results support CIN as a promising potential biomarker of radiation response, which would discriminate patients eligible for dose de-escalation from those requiring additional radiosensitizers such as docetaxel.

## Results

### Radiation induces CIN in head and neck cancer

We first characterized and quantified the CIN induced by 2 Gy of IR, which represents the most common daily dose used to treat head and neck cancer patients, in three HPV-negative and four HPV-positive head and neck cancer cell lines. 2 Gy significantly increased levels of misaligned chromosomes, lagging chromosomes and chromosome bridges after 24 hours (Fig. 1A-D). Radiation has previously been reported to induce multipolar spindles at high doses (64,65) which was found to be due to radiation-induced centrosome amplification (66–68). However, 2 Gy increased multipolar spindles only in the SCC-22B HPV-negative cell line (Fig. 1E). In total, radiation increased the incidence of abnormal mitotic figures consistent with CIN an average of 39% in early stages of mitosis and 56% in late stages of mitosis 24 hours after 2 Gy (Fig. 1F).

**Figure 1.**
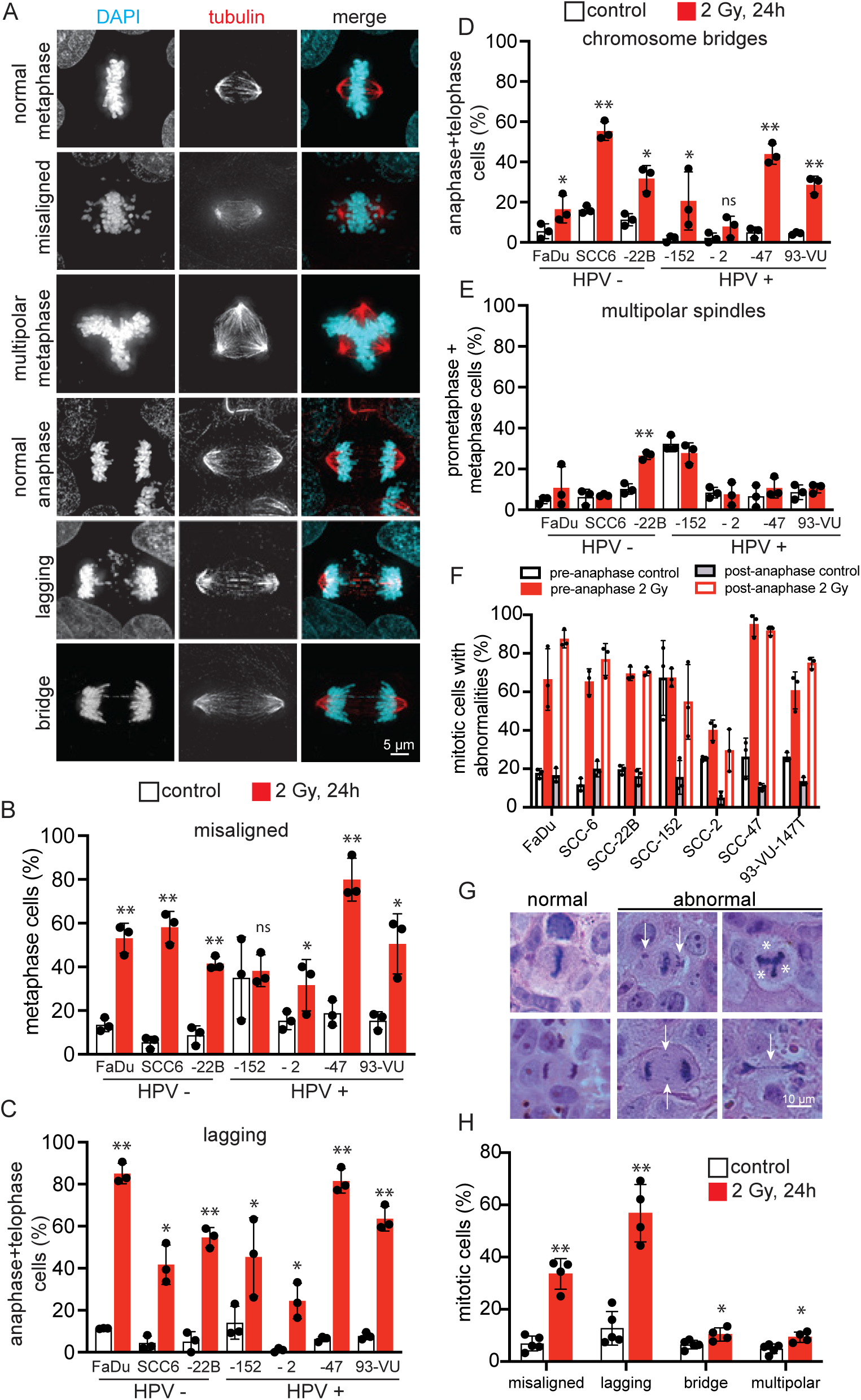
Radiation induces multiple types of CIN in head and neck cancer cells. (A) Representative images of types of chromosome segregation errors that lead to CIN in FaDu head and neck cancer cells 24 hours after 2 Gy of radiation. Normal metaphase and anaphase are shown for reference. (B-E) Quantification of different types of mitotic defects that cause CIN in HPV-negative (HPV-) and HPV-positive (HPV+) untreated control cells and 24 hours after 2 Gy of radiation. In (B), misaligned chromosomes include polar chromosomes. n≥50 cells in each phase of mitosis (prometaphase, metaphase, anaphase, and telophase) per condition in each of 3 biological replicates. (F) Quantification of the total CIN in each cell type before and 24 hours after 2 Gy of radiation. Misaligned chromosomes and multipolar spindles contribute to pre-anaphase CIN, while lagging and bridge chromosomes contribute to post-anaphase CIN. n=50 cells per condition in each of 3 biological replicates. (G) Representative H&E images of FaDu tumor xenografts 24 hours after 2 Gy radiation. Upper panel shows metaphase cells. From left to right: normal, misaligned chromosomes (arrows), multipolar spindle (*denotes inferred spindle pole). Lower panel shows anaphase cells. From left to right: normal, lagging chromosome (arrows), chromosome bridge (arrow). (H) Quantification of radiation-induced CIN in vivo. An average of 79 metaphases (range 36-132) and 35 anaphase/telophase (range 13-71) were counted per tumor. n=4 tumors in each condition. Error bars indicate SD. Statistical differences determined by 2-tailed t-test, * = p<0.05, ** = p<0.001.

We then quantified the percentage of misaligned and lagging chromosomes that contained centromeres as an indication of the ratio of whole chromosome to structural CIN induced by IR. Over 85% of misaligned and 90% of lagging chromosomes after radiation lacked a centromere (Supp Fig. S1), consistent with structural CIN induced by unrepaired, or erroneously repaired, double-stranded DNA breaks. These acentric fragments do not contain a kinetochore and are therefore unable to attach to microtubules, leading to their missegregation. Thus, the conventional dose of radiation in head and neck cancer patients induces primarily structural but also numerical CIN in head and neck cancer cells.

We then quantified radiation-induced CIN in a mouse tumor model. HPV-negative FaDu murine xenografts were treated with sham radiation or 2 Gy and tumors were harvested and fixed 24 hours after radiation. Analysis of H&E-stained slides revealed a substantial increase in misaligned and lagging chromosomes 24 hours after 2 Gy, with little increase in chromosome bridges and multipolar spindles (Fig. 1G-H). These findings were corroborated in an independent cohort of murine xenografts composed of one HPV-negative and three HPV-positive cell lines (Supp Fig. S2). 2 Gy of ionizing radiation substantially increased misaligned and lagging chromosomes with only modest effects on chromosome bridges and multipolar spindles (Supp Fig. S2). Together, these results indicate that the daily dose of radiation commonly given to patients causes substantial CIN due to misaligned and lagging chromosomes in HPV-positive and HPV-negative head and neck cancer cells in vitro and in vivo.

### Chromosomal instability sensitizes cells to radiation

Combining two tolerable sources of CIN increases CIN over the maximally tolerated threshold leading to cell death (22,35,36,42). We therefore hypothesized that cells with higher levels of CIN at baseline would be more sensitive to radiation, since radiation induces CIN. To test this hypothesis, we induced CIN in FaDu HPV-negative head and neck cancer cells by knocking down the mitotic spindle assembly checkpoint protein Mad1 (Mitotic Arrest Deficient 1) using shRNA (Fig. 2A). Mad1 recruits its binding partner Mad2 to unattached kinetochores, where Mad2 is converted into an active inhibitor of the Anaphase Promoting Complex/Cyclosome (6,33,69). Though Mad1 is essential, cells survive partial depletion but show weakened mitotic checkpoint activity and CIN (33). Mad1 overexpression also weakens mitotic checkpoint signaling by sequestering Mad2 in the cytoplasm leading to CIN (6). Increased or decreased expression of the gene encoding Mad1, MAD1L1, occurs in 27% of head and neck cancers (Supp Fig. S3). As expected, ∼50% knockdown of Mad1 increased lagging and bridge chromosomes, consistent with CIN (Fig. 2A-C). Importantly, standard clonogenic assays revealed that increasing CIN by Mad1 knockdown increased radiosensitivity in head and neck cancer cells (Fig. 2D). Since mitosis is the most radiosensitive stage of the cell cycle (70), we tested whether the increased radiosensitivity caused by Mad1 knockdown could be due to an increase in the percentage of mitotic cells. However, reduced expression of Mad1 did not affect mitotic index (Supp Fig. S4A).

**Figure 2.**
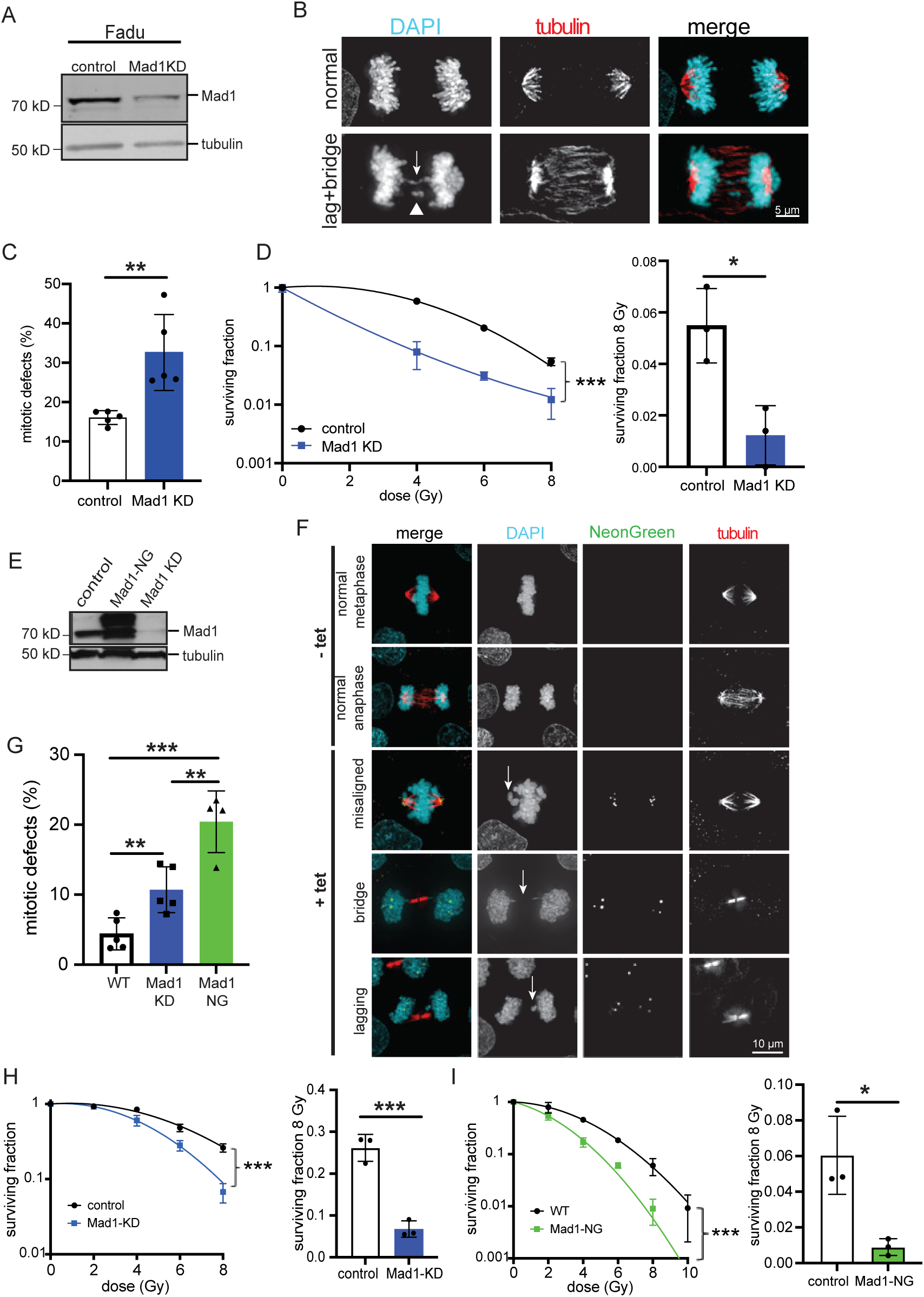
CIN sensitizes to radiation in head and neck and cervical cancer cells. (A-D) Mad1 knockdown (KD) induces CIN in FaDu HPV-negative head and neck cancer cells and sensitizes them to radiation. (A) Immunoblot showing efficient Mad1 knockdown. (B) Images of (top) normal anaphase and (bottom) anaphase cell with chromosome bridge (arrow) and lagging chromosome (arrowhead). (C) Quantification of mitotic defects (lagging and bridge chromosomes) in isogenic parental and Mad1 knockdown FaDu cells. n≥60 cells in metaphase and ≥70 cells in anaphase+telophase per condition in each biological replicate. (D) Clonogenic assays showing Mad1 knockdown FaDu cells with CIN have increased radiation sensitivity relative to isogenic parental cells. (E-I) Induction of CIN in HeLa HPV-positive cervical cancer cells sensitizes them to radiation. (E) Immunoblot showing expression of Mad1-mNeonGreen (NG) and Mad1 knockdown in HeLa cells. (F) Images showing examples of normal and abnormal mitotic figures in HeLa cells +/- tet inducible expression of Mad1-NG. Arrows indicate respective defect. (G) Quantification of mitotic errors due to Mad1 knockdown and expression of Mad1-NG, which both cause CIN. n≥70 cells in metaphase and ≥85 cells in anaphase or telophase per condition in each biological replicate. (H) Clonogenic assays showing that both Mad1 knockdown and Mad1-NG expression sensitize cells to radiation. The surviving fraction of cells after 8 Gy is shown to the right of each respective clonogenic curve. n=3 biological replicates each. Error bars indicate SD. Statistical differences determined by 2-tailed t-test, * = p<0.05, ** = p<0.01, *** = p<0.001.

To test whether this effect was specific to head and neck cancer or conserved in other cancer types, we extended this analysis to cervical cancer cells. Alterations in Mad1 expression are as common in cervical cancer as in head and neck cancer (Supp Fig. S3). We therefore increased CIN in HeLa cervical cancer cells using two methods: Mad1 knockdown and tetracycline-inducible overexpression of Mad1. As expected, stable Mad1 knockdown and tetracycline-inducible expression of Mad1-mNeonGreen (NG) both caused an increase in lagging and bridge chromosomes associated with CIN (Fig. 2E-G). Neither Mad1 knockdown nor expression of Mad1-NG affected cell cycle timing or mitotic index (Supp Fig. S4B-C). We then tested whether increasing CIN enhanced radiosensitivity in these isogenic cell lines using clonogenic assays. As hypothesized, both CIN cell lines were significantly more radiosensitive than their parental isogenic counterparts (Fig. 2H-I). Together, these results demonstrate that baseline CIN sensitizes otherwise isogenic head and neck and cervical cancer cells to radiation.

### Docetaxel causes cell death by inducing CIN due to multipolar spindles without mitotic arrest

Docetaxel is a well-known radiosensitizer that increased median overall survival from 15.3 to 25.5 months when added to radiation therapy in a recent phase III trial of head and neck cancer patients (58). Docetaxel, like paclitaxel, has been thought to increase radiation sensitivity by increasing the percentage of cells in mitosis (50–52). However, based on the recent findings with paclitaxel in breast cancer (20–22,63), we tested the hypothesis that docetaxel radiosensitizes because it induces multipolar spindles rather than mitotic arrest in head and neck cancer. HPV-negative FaDu cells were treated with a range of low doses of docetaxel and the incidence of multipolar spindles during prometaphase and metaphase (pre-anaphase) and anaphase and telophase (post-anaphase) was quantified. Like paclitaxel, docetaxel induced an increase in both pre- and post-anaphase multipolarity in a concentration dependent manner (Fig. 3A-B). At each concentration, the incidence of multipolar spindles was higher at early stages of mitosis than at later stages, due to cells focusing multipolar spindles into bipolar spindles, as previously shown for paclitaxel (20–22). Cells that focus multipolar spindles into bipolar spindles early in mitosis produce daughter cells that generally survive, while those that maintain multipolar spindles throughout mitosis typically produce three daughter cells that are inviable (9,20–22). Thus, the incidence of multipolar spindles that persist into late stages of mitosis is more predictive of cell fate. Importantly, cells in later stages of mitosis were readily detectable in each of these concentrations of docetaxel, which is inconsistent with mitotic arrest, in which cells arrest in prometaphase. Docetaxel concentrations ≥0.3 nM induced multipolar divisions in >20% of cells, while multipolar spindles persisted into late stages of anaphase in <10% of cells treated with lower concentrations. At least 10-fold higher concentrations of docetaxel were necessary to induce mitotic arrest (Fig. 3C). The anti-proliferative activity of docetaxel correlated closely with multipolar spindle polarity in late mitosis, with docetaxel concentrations ≥0.3 nM inducing significantly decreased viability, while lower concentrations had minimal effect (Fig. 3D). Similar results were found with the HPV-positive cell line SCC-47 (Supp Fig. S5), indicating that this mechanism occurs independently of HPV status. As in HPV-negative FaDu cells, the anti-proliferative effects of docetaxel closely correlated with its ability to induce multipolar spindles that persisted into late stages of mitosis in >20% of cells (Supp Fig. S5B-C). Again, these effects occurred at concentrations ≤10-fold lower than those necessary to induce mitotic arrest (Supp Fig. S5D). Thus, low doses of docetaxel induce cell death via formation of multipolar spindles without causing mitotic arrest in both HPV+ and HPV- head and neck cancer cells.

**Figure 3.**
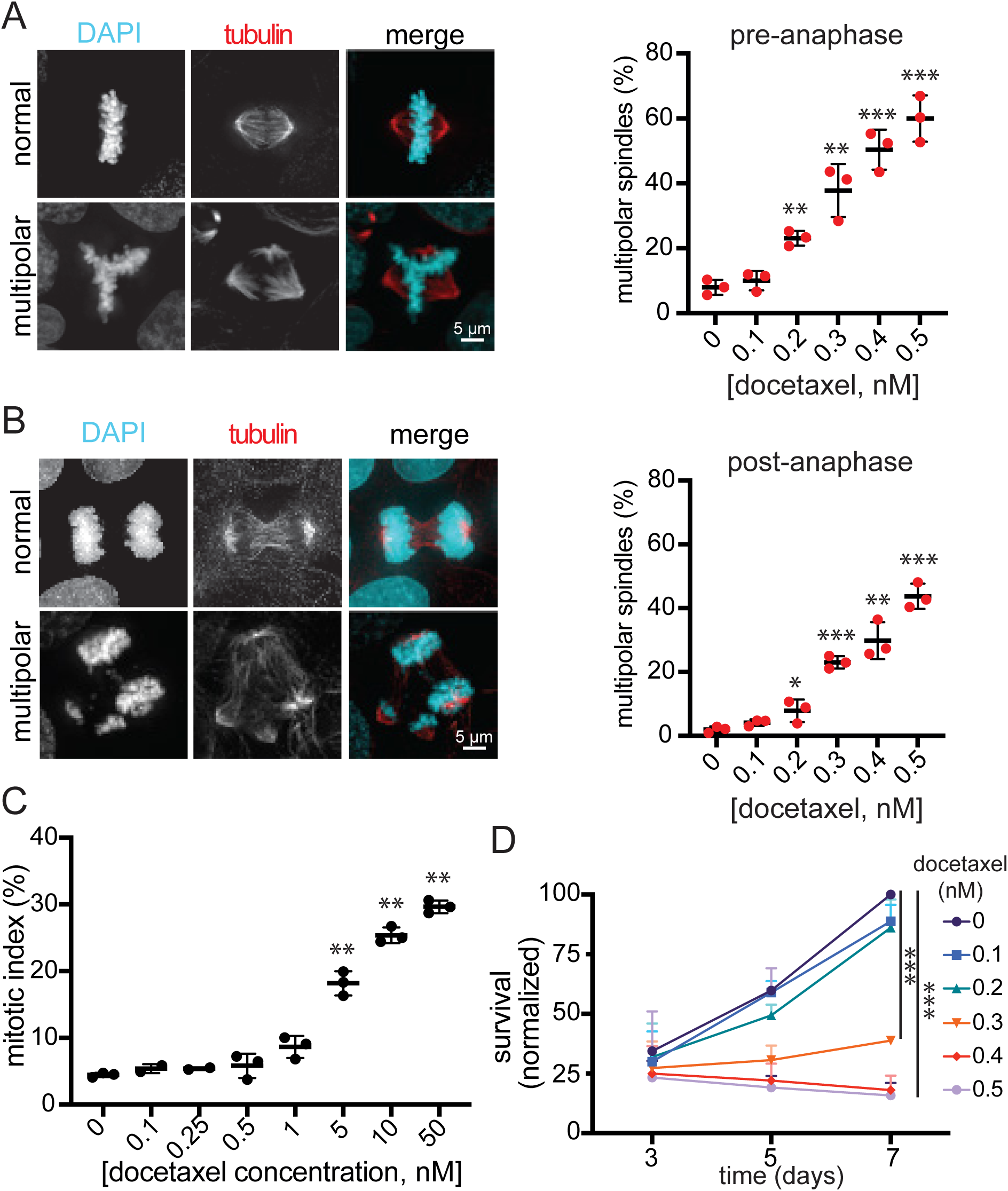
Docetaxel causes cell death in head and neck cancer cells by inducing CIN on multipolar spindles without mitotic arrest. (A) Left: Images of FaDu cells in metaphase showing normal bipolar spindle (top) or abnormal multipolar spindle (bottom). Right: Quantification of pre-anaphase multipolar spindles (in prometaphase and metaphase cells), which increase in a concentration dependent manner 24 hours after treatment with docetaxel. (B) Left: Images of FaDu cells in normal bipolar anaphase (top) and abnormal multipolar anaphase (bottom). Right: Quantification of multipolar spindles in post-anaphase (anaphase and telophase) cells 24 hours after docetaxel treatment. (A-B) n=100 cells per condition in each of 3 biological replicates. (C) Mitotic index after 24 hour treatment with the indicated concentrations of docetaxel showing that the concentrations used in A and B that induce multipolar spindles are too low to cause mitotic arrest. n≥500 cells in each of 3 biological replicates. (D) MTT assay showing concentrations of docetaxel that induce ≥20% multipolar spindles in post-anaphase cells impair proliferation. Y-axis values normalized to DMSO-treated cells at day 7. n=3 biological replicates. Error bars indicate SD. Statistical differences determined by 2-tailed t-test, * = p<0.05, ** = p<0.01, *** = p<0.001 versus DMSO.

We questioned whether the well-known radiation sensitizing effects of docetaxel are due to mitotic arrest, as long expected, or CIN due to division on multipolar spindles. FaDu and SCC-47 cells were treated with 0.2 nM docetaxel, a concentration of docetaxel that induced low levels of multipolar spindles but not cell death or mitotic arrest, for 24 hours followed by irradiation. Clonogenic assays revealed this low level of docetaxel increased the radiosensitivity of both HPV- and HPV+ cell lines (Fig. 4A-B).

**Figure 4.**
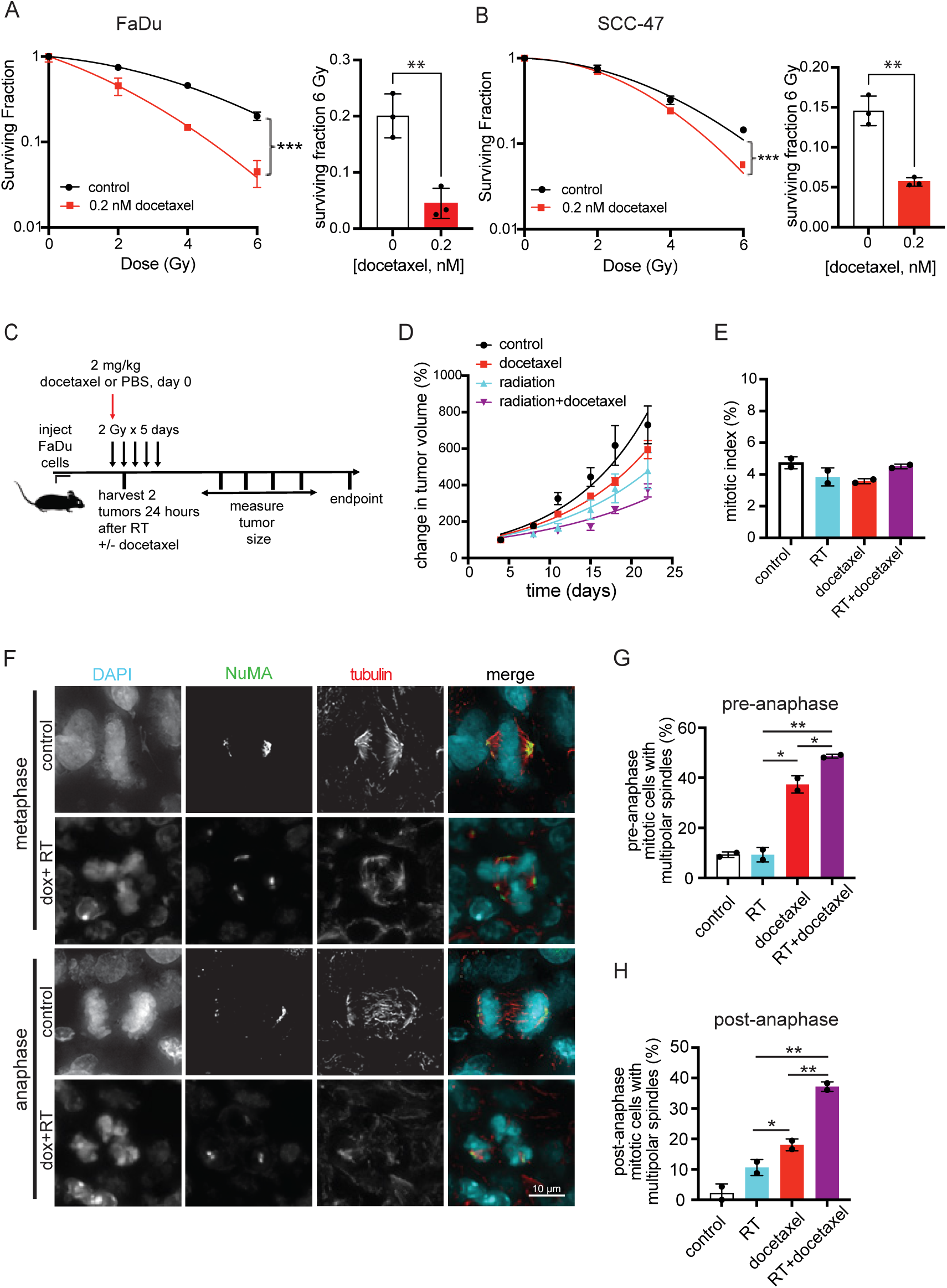
Docetaxel sensitizes HPV-positive and HPV-negative head and neck cancer cells to radiation by inducing CIN on multipolar spindles without mitotic arrest. (A-B) Clonogenic assays showing docetaxel concentrations that are too low to cause mitotic arrest (0.2 nM) sensitize HPV-negative FaDu (A) and HPV-positive SCC-47 (B) cells to radiation. The surviving fraction of cells after 6 Gy is shown to the right of each respective clonogenic curve. n=3 biological replicates. (C) Schematic of murine study. RT = radiation therapy. (D) Docetaxel sensitizes FaDu xenograft tumors to radiation. n=10 tumors per condition. Error bar, SEM. (E) Mitotic index in murine tumors is not elevated relative to PBS control by treatment with docetaxel, radiation, or docetaxel+radiation. n≥500 cells in each of 2 tumors per condition. (F-H) Docetaxel induces multipolar spindles in FaDu xenograft tumors. (F) Immunofluorescence images of normal bipolar and abnormal multipolar spindles in FaDu xenograft tumors. Spindle poles were identified by co-localization of NuMA and α-tubulin. (G-H) Quantification of multipolar spindles in xenograft tumor tissue in (G) pre-anaphase cells (prometaphase and metaphase) and (H) post-anaphase cells (in anaphase and telophase). n≥80 (range 83-187) pre-anaphase and ≥23 (range 23-57) post-anaphase cells in each of 2 biological replicates. Error bars indicate SD. Statistical differences determined by 2-tailed t-test * = p<0.05, ** = p<0.01.

To determine if this novel mechanism of radiosensitization occurs in vivo, we injected athymic nude mice harboring FaDu tumors with PBS or a low dose of docetaxel. Two hours later we irradiated the tumors with 2 Gy. Daily irradiation with 2 Gy continued for an additional 4 days, mimicking patient treatment schedules, with sham irradiation as a control (Fig. 4C). Two tumors from each group of mice were harvested 24 hours after their respective treatment and pre- and post-anaphase multipolar spindles and mitotic index were quantified in each tumor. Both nuclear mitotic apparatus protein (NuMA) and α-tubulin were used as spindle pole markers. As expected, docetaxel decreased tumor cell growth in combination with radiation compared to treatment with docetaxel or radiation alone (Fig. 4D). Inclusion of docetaxel substantially increased the incidence of multipolar spindles after radiation without affecting mitotic index (Fig. 4E-H). Importantly, docetaxel treatment increased multipolar spindles in late stages of mitosis from approximately 10% in tumors treated with radiation alone to almost 40% in combination with radiation (Fig. 4H). These data support the hypothesis that docetaxel sensitizes head and neck tumors to radiation by inducing multipolar spindles that persist into anaphase and cause CIN, rather than by causing mitotic arrest.

### CIN directly correlates with increased radiation response in HPV+ and HPV- patient-derived xenografts treated with definitive radiation

To determine if CIN is associated with increased radiation response in tumors from head and neck cancer patients, we first used well-established HPV+ and HPV- head and neck cancer patient-derived xenograft (PDX) tumors (71). Patient-derived tumor tissues (4 HPV-positive and 5 HPV-negative) were each grown in athymic nude mice (12-14 tumors per treatment group). To establish baseline rates of CIN, mitotic defects were quantified in the tumors of untreated mice using immunofluorescence microscopy (Fig. 5A). Tumor size 2 weeks following 2 Gy radiotherapy daily for 5 days was compared in the treated versus sham-treated mice to quantify tumor regression. In 8/9 cases, the correlation between CIN and response to radiation clustered according to HPV status (Fig. 5B). Within tumors of a given HPV status, CIN directly correlated with the extent of tumor regression in response to radiotherapy (Fig. 5B), supporting the conclusion that tumors with higher levels of CIN prior to treatment are more sensitive to radiation and suggesting that HPV reduces the maximally tolerated threshold of chromosome loss.

**Figure 5.**
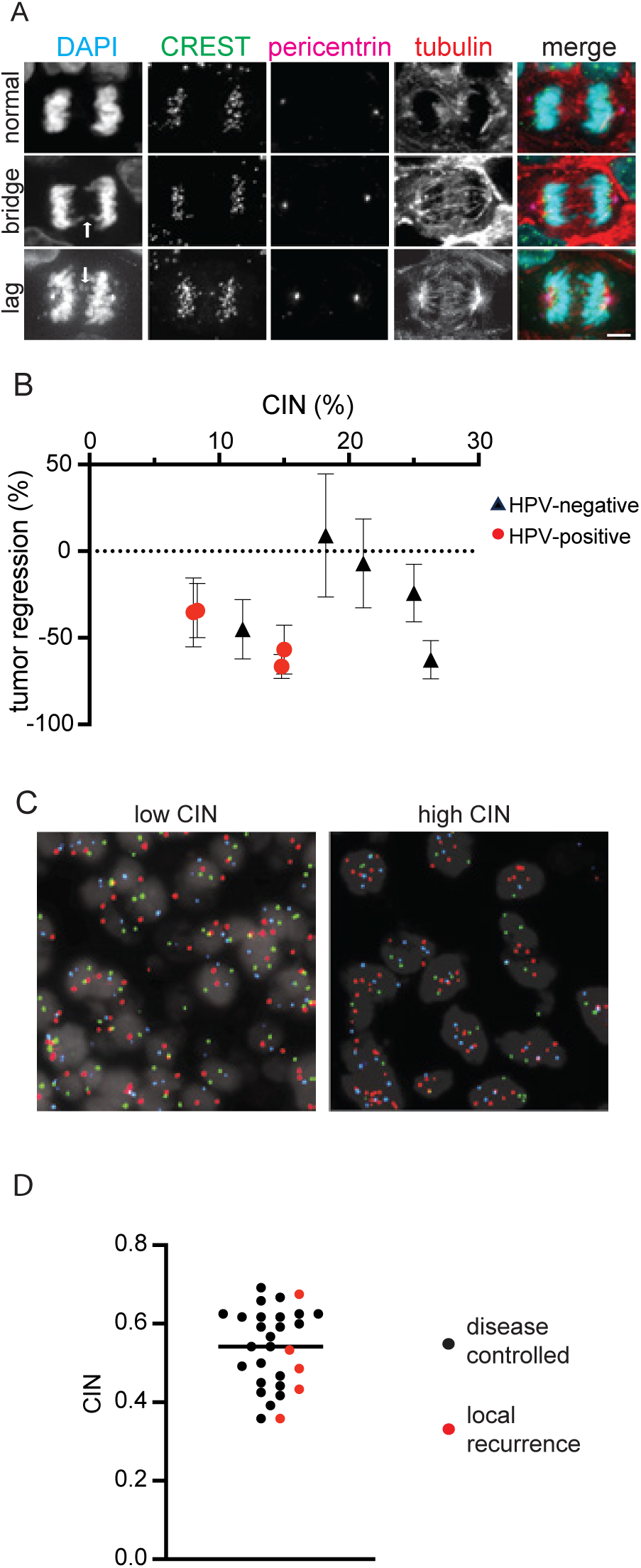
CIN directly correlates with radiation response in head and neck cancer PDX tumors and laryngeal cancer patients. (A) Images of normal anaphase and anaphase defects in head and neck cancer PDX tissues prior to irradiation. Arrows indicate the indicated defect. Scale bar, 5 μm. (B) PDX tumors with higher CIN are more sensitive to radiation. CIN is reported as the percentage of anaphase and telophase cells containing lagging chromosomes and/or chromosome bridges. Tumor regression is based on 12-14 tumors per PDX and treatment group treated with sham or 2 Gy daily for 5 days. Error bars represent SD. n≥19 cells in anaphase/telophase (range = 19-139, average = 47) per sample to quantify anaphase defects. (C) Interphase FISH images used to quantify CIN in laryngeal cancer. Two sections with centromeric probes (CEP) to chromosomes 3, 7, 9 (left, red, blue, green respectively) and 4, 10, 17 (right: green, blue, red, respectively) were used to quantify CIN for each sample. (D) Laryngeal tumors with CIN below the median (black bar, 0.54) had increased local recurrence rate (31%) compared to tumors with CIN above the median (6%).

### Increased baseline CIN associates with improved response in laryngeal tumors treated with definitive radiation

To determine if CIN impacts sensitivity of head and neck cancers to radiation in the clinical setting, we identified 29 patients with locally advanced laryngeal cancer who were treated with definitive radiation therapy (with or without chemotherapy) and for whom biopsy tissue and follow-up data were available. 18/29 (62%) patients received chemotherapy concurrently with radiation and the other 38% were treated with radiation alone. Patients who received surgery were excluded from this study. Patient and treatment characteristics are detailed in Table 1. The vast majority (94-100%) of laryngeal cancers are not associated with HPV (72–74), thus most samples were not tested for p16. The small size of the diagnostic cancer biopsies prohibited quantification of CIN by scoring aberrant mitotic figures, since there was an insufficient number of mitotic cells in each tumor sample for accurate quantification. We therefore quantified CIN based on another well-established method (22,44,45,75) that has been shown to correlate well with direct quantification of mitotic errors (25,76), intercellular variability in the copy number of 6 chromosomes using interphase centromere FISH (Fig. 5C). CIN is scored as the percentage of cells that deviate from the modal copy number for each of the 6 chromosomes, averaged over the 6 chromosomes, and thus represents a measure of the variability of chromosome numbers between cells. Normal tonsillar epithelia obtained from benign tonsillectomy samples were used as normal control tissue for CIN. The range of CIN in the tumor tissues was 36 – 69%, indicating significant heterogeneity in terms of chromosome missegegration rates between patients. CIN in tumors was higher than control tissue (Supp Fig. S6), consistent with what has been found in normal and cancerous breast tissue using the same method (25). With a median follow-up of 4.5 years, five patients (17%) either did not respond to RT completely as evidenced by residual viable tumor following radiotherapy, or had recurrent disease in the irradiated field indicating radiation resistance. Interestingly, 4/5 of the patients who recurred locally had CIN below the median of the cohort (Fig. 5D), consistent with insufficient rates of baseline CIN conferring resistance to radiation. Thus, laryngeal cancer patients with CIN at or below the median of the cohort had a 31% rate of recurrence, while patients with CIN above the median only had a 6% rate of recurrence. Together, these results support the hypothesis that head and neck cancer cells with higher levels of CIN at baseline are more sensitive to radiation, as indicated by decreased local recurrence.

**Table 1.** Patient characteristics of the University of Wisconsin laryngeal cancer cohort. Disease site (AEF, aryepiglottic fold; TVC, true vocal cord; AC, anterior commissure). p16 status is a surrogate for HPV positivity. NK is not known/not tested. *denotes a patient that experienced disease recurrence/incomplete response to radiation.

| Patient | Sex | Age | Disease site | Stage | Radiation dose (Gy) | Chemo-therapy | p16 |
| --- | --- | --- | --- | --- | --- | --- | --- |
| 1* | M | 58 | Epiglottis | T3N2cM0 (IV) | 72 | none | NK |
| 2 | M | 50 | Epiglottis/AEF | T3N0M0 (III) | 72 | none | NK |
| 3* | F | 51 | AEF | T3N0M0 (III) | 70.2 | cisplatin | NK |
| 4 | M | 47 | Epiglottis | T2N3M0 (IV) | 73.2 | cisplatin | NK |
| 5 | M | 66 | Epiglottis/AEF | T3N1M0 (III) | 70 | cisplatin | NK |
| 6* | M | 60 | Subglottis/TVC | T3N0M0 (III) | 72 | none | NK |
| 7 | F | 60 | Epiglottis | T3N0M0 (III) | 69.6 | cisplatin | NK |
| 8 | M | 82 | TVC | T1N0M0 (I) | 63 | none | NK |
| 9* | M | 56 | TVC | T4N2cM0 (IV) | 74.8 | unknown | NK |
| 10 | M | 45 | Epiglottis | T3N0M0 (III) | 69.6 | cisplatin | NK |
| 11 | F | 64 | Supraglottis | T4N2cM0 (IV) | 70 | cetuximab | NK |
| 12 | M | 62 | Supraglottis | T3N2cM0 (IV) | 70 | carboplatin | NK |
| 13 | M | 65 | TVC | T2N0M0 (II) | 65.25 | none | NK |
| 14 | M | 68 | Larynx | T3N2bM0 (IV) | 70 | cisplatin | NK |
| 15 | M | 64 | Epiglottis | T3N2bM0 (IV) | 70 | unknown | NK |
| 16* | M | 61 | Supraglottis | T4aN2bM0 (IV) | 70 | cetuximab | + |
| 17 | F | 58 | Supraglottis | T3N0M0 (III) | 70 | cisplatin | NK |
| 18 | M | 62 | TVC, epiglottis | T3N0M0 (III) | 70 | none | NK |
| 19 | M | 53 | Supraglottis | T3N2cM0 (IV) | 72 | cisplatin | - |
| 20 | M | 57 | TVC | T3N1M0 (III) | 63.63 | cetuximab | NK |
| 21 | M | 69 | Supraglottis | T2N2cM0 (IVA) | 70 | cisplatin | NK |
| 22 | M | 83 | TVC | T3N0M0 (III) | 70 | none | NK |
| 23 | M | 64 | Supraglottis | T4aN0M0 (IV) | 70 | cisplatin | - |
| 24 | M | 29 | TVC | T2N0M0 (II) | 72.8 | none | + |
| 25 | M | 62 | Supraglottis | T3N2cM0 (IV) | 70 | cisplatin | - |
| 26 | M | 72 | TVC | T2N0M0 (II) | 65.25 | none | NK |
| 27 | M | 67 | AC, TVC, ventricle, false cord | T3N0M0 (III) | 70 | none | NK |
| 28 | F | 76 | AEF | T2N0M0 (II) | 70 | none | NK |
| 29 | F | 54 | AEF | T1N1M0 (III) | 70 | cisplatin | NK |

## Discussion

We have shown in two tissue contexts that otherwise isogenic cancer cells with higher levels of CIN at baseline are more sensitive to radiation, likely because they are closer to their maximally tolerated threshold of chromosome loss. Head and neck cancer PDX tumors with increasing levels of baseline CIN are more sensitive to definitive radiation, with HPV-positive PDXs being more sensitive overall than HPV-negative tumors, consistent with clinical findings. Advanced laryngeal cancers with baseline CIN above the median value had a substantially lower rate of recurrence than those with CIN below the median (6% vs 31%). Overall, these results demonstrate that higher baseline CIN sensitizes cells to radiation in head and neck and cervical cancer cell culture models as well as head and neck PDX and patient tumors, providing a foundation for CIN as a biomarker of radiation response. Such a biomarker could have a profound impact on treatment strategies by permitting dose de-escalation with reduced toxicity in patients with radiosensitive tumors, as well as more aggressive therapy in patients with radioresistant cancers.

Docetaxel has been used as a radiation sensitizer for almost half a century and has improved survival in head and neck cancer patients (54,58). The radiosensitizing effect of docetaxel was recently corroborated in a phase III randomized-controlled trial comparing radiation alone to docetaxel with radiation in patients with locally advanced head and neck cancer (58). It has long been accepted that docetaxel acts as a radiosensitizer by arresting cells in mitosis, the most radiosensitive stage of the cell cycle. However, we recently showed that concentrations of paclitaxel in primary breast cancers are too low to cause mitotic arrest (22,63). Instead, clinically relevant concentrations of paclitaxel induce multipolar divisions and increase CIN. In a cell-based assay, the addition of paclitaxel to radiation substantially increased CIN, assessed by missegregation of a human artificial chromosome, and had synergistic effects on cell death, suggesting that paclitaxel sensitizes to radiation by increasing CIN (77). Here, we show for the first time that docetaxel-induced radiosensitization can occur without mitotic arrest, which is contrary to the presumed mechanism over the last 40 years. Rather, docetaxel can sensitize to radiation by inducing CIN in the form of multipolar spindles that persist throughout mitosis. In the presence of CIN-inducing radiation, low doses of docetaxel cause cell death by increasing CIN over a maximally tolerated threshold. Though we hypothesize this to be the mechanism of radiosensitization in head and neck cancer patients, intratumoral concentrations of docetaxel have not been quantified in this patient population, and the docetaxel concentrations used herein have been extrapolated from similar studies in breast tumors (22,63). We are currently planning a clinical trial of head and neck cancer patients analogous to Scribano et al. (22), to determine if docetaxel induces multipolar spindles in patient tumors.

HPV-positive tumors are generally more responsive to chemoradiation than their HPV-negative counterparts. In the head and neck cancer context, patients with HPV- positive tumors have a significantly better prognosis than patients with HPV-negative cancers (47). Though we have previously shown that HPV+ head and neck cancer cells and tumors have more of a specific type of CIN (polar chromosomes) than HPV- tumors, total CIN does not differ based on HPV status (78). This implies that the enhanced radiation sensitivity clinically observed in HPV+ tumors could be attributed to factors other than CIN, such as low levels of active p53, prolonged and impaired DNA damage repair, or decreased hypoxia (79–81). However, our PDX data suggest that HPV-positivity lowers the maximally tolerated CIN threshold, perhaps due to the presence of low levels of wild-type p53, which detects structural aneuploidy and CIN. Overall, our results imply that within HPV+ or HPV- cancers, tumor cells with high CIN are likely to be more radiosensitive.

Our results demonstrating that increasing baseline CIN sensitizes to radiation are consistent with prior work showing that reducing CIN in glioma cells and orthotopic tumors causes resistance to radiation (12). Similarly, higher rates of CIN (based on cell-to-cell copy number variability in single cell sequencing data) were associated with radiation sensitivity in colorectal cancer organoids (82). Increasing CIN above a maximally tolerated threshold can be achieved in multiple ways (4,14,83). Recently, depleting a mitotic kinase that controls genome stability, MASTL, was shown to increase CIN in prostate cancer cells, leading to cell death (42). Pharmacologically, inhibition of TTK protein kinase (also known as monopolar spindles 1, MPS1) sensitized breast cancer cells to radiation (84). This was reportedly due to decreased efficiency of homologous recombination, though Mps1 inhibition is commonly used to increase mitotic errors (85), and increased CIN was likely involved in the enhanced radiosensitivity observed. While increased CIN can enhance radiation sensitivity, it is not known if one particular type of CIN confers this advantage over another. It is likely that the type of CIN is less important than the rate of CIN.

Radiation can directly induce double stranded DNA breaks, which if left unrepaired or repaired erroneously, can cause CIN. We have shown in head and neck cancer cells that approximately 90% of missegregated chromosomes induced by 2 Gy of radiation are acentric fragments. This demonstrates that head and neck cancer cells proceed into mitosis despite significant unrepaired DNA damage, implying an impaired DNA damage checkpoint. p53 is essential for cell cycle arrest or apoptosis in the presence of DNA damage. *TP53* is mutated in 86% of HPV-negative head and neck cancers (86). In HPV-positive cancers, p53 is degraded by the HPV oncoprotein E6 (87), rendering these cells p53 deficient. Thus, absence of functional p53 may allow these cells to proceed into mitosis in the presence of radiation-induced DNA damage, resulting in structural CIN. Our results demonstrate that an impaired DNA damage response can be exploited to promote tumor cell death by increasing CIN above a maximally tolerated threshold.

6-centromere FISH can reliably quantify CIN using an assay that is routine in clinical labs worldwide. However, this approach is limited by its ability to detect copy numbers of only 6 centromeres, and future methods that can detect copy numbers of all chromosomes and chromosome arms may further improve quantification of CIN in clinical samples. Importantly however, bulk sequencing and transcriptome analysis, which can measure recurrent aneuploidy that is shared among a large percentage of the cell population, are unable to detect ongoing CIN (76). Advances in single cell sequencing, which is superior to bulk sequencing for the detection of CIN, may further improve CIN measures in clinical samples in the future. In the meantime, identification of a CIN threshold that increases radiation response based on 6-centromere FISH analysis is a critical step in translating this potential biomarker to clinic.

Currently, multiple clinical trials are testing the impact of reducing radiation dose in HPV-positive head and neck cancers given their overall improved outcomes. However, most are not using specific biological variables to further categorize patients into specific risk groups. Our data show that tumors with higher CIN are more sensitive to radiation. Following further validation, these results could provide a basis for the use of CIN as a biomarker for selection of patients eligible for dose de-escalation, which could thereby improve patient outcomes by reducing side effects that negatively impact quality of life. An even greater clinical need is to improve response for patients with HPV-negative head and neck cancer, given that only 56% of these patients are free of disease at 5 years (88). Patient tumors with lower-than-average baseline CIN may be less likely to respond to radiation, and this could be an indicator that addition of a CIN-inducing drug would enhance radiation response by increasing CIN above a maximally tolerated threshold. Taxane therapy with docetaxel or paclitaxel, inhibition of the mitotic kinesin CENP-E, and inhibition of the mitotic checkpoint kinase Mps1 all have the potential to elicit this effect (21,22,85,89,90) and could provide novel strategies for enhancing radiosensitivity in head and neck cancer.

## Methods

### Cell lines and treatments

All head and neck and cervical cell lines were confirmed via short tandem repeat testing and frequently tested for mycoplasma. Head and neck cancer cell lines include: FaDu (HPV-), UM-SCC-6 (HPV-), UM-SCC-22B (HPV-), UPCI SCC-152 (HPV16+), UM-SCC-2 (HPV16+), UM-SCC-47 (HPV16+), 93-VU-147T (HPV16+). Cervical cell line: HeLa (HPV18+). All cells were grown in DMEM with 4.5 g/dL glucose, 10% FBS, and 1% penicillin/streptomycin, with the exception of SCC-152 cells, which were grown in MEM with 10% FBS and 1% pen/strep. Each line was maintained at 37°C and 5% CO_2_.

Cells were plated on coverslips, allowed to adhere for 24 hours, and then irradiated using an Xstrahl X-ray System, Model RS225 (Xstrahl, UK) at a dose rate of 3.27 Gy/min at 30 cm FSD, tube voltage of 195 kV, current of 10 mA and filtration with 3 mm Aluminum. Docetaxel (Sigma) was diluted in DMSO to a stock solution of 12.4 μM and stored at −20°C for up to 3 months.

HeLa-Mad1 KD cells were described previously (91). To create the Mad1 overexpressing cells, HeLa cells stably expressing the tet repressor (91) were transduced with retrovirus expressing full length wild type Mad1 tagged with mNeonGreen at the C-terminus under a tet-inducible promoter. Stable integrants were selected with 4 μg/mL puromycin (Mad1-mNeonGreen) and 200 µg/mL blasticidin (tet) and validated for inducible expression of Mad1 upon tet addition. To obtain subclones with uniform Mad1-NG expression, the top 30% of NG-positive cells were collected using a flow cytometer and plated immediately for clonogenic assay. To create FaDu Mad1-KD cells, retrovirus expressing shRNA against Mad1 (5′-AGCGATTGTGAAGAACATG-3′) was made from pSUPERIOR.retro.puro as in (91). FaDu cells with stable integration of Mad1-KD shRNA were selected with 2 μg/mL puromycin. These parental cells were subcloned and a clone with at least 50% knockdown of Mad1 as verified by Western blot was selected for further experiments. All Mad1 overexpressing or KD cell lines were kept under puromycin selection.

### MTT proliferation assays

Cell survival and proliferation was quantified using MTT assay. Briefly, cells were counted and plated in 6 well plates and treated the next day. On the day(s) of measurement, cells were incubated with 1 mg/mL MTT reagent (3-(4,5 dimethylthiazol-2-yl)-2,5diphenyltetrazolium bromide) for 3 hours. Formazan, the metabolic end product measured in this assay, was released from cells by adding 800 μL of DMSO and 100 μL Sorenson’s glycine buffer (0.1M glycine, 0.1M NaCl, pH 10.5 with 0.1M NaOH) to each well, followed by a 10 min incubation at 37°C. Formazan was detected by measuring absorbance on a plate reader at 540 nm.

### Clonogenic assays

Cells were trypsinized, counted, plated into 6 well plates, allowed to adhere, and then irradiated using the Xstrahl X-ray System as above. In the clonogenic assays with docetaxel, cells were plated, treated with 0.2 nM docetaxel for 24 hours, and then irradiated. Cells were allowed to grow for 10-14 days and then fixed with pure methanol for 20 minutes and stained with 1% crystal violet. Colonies containing at least 50 cells were counted under a dissection microscope. Each experiment was performed in triplicate and was repeated at least 3 times. Survival curves were compared using a non-linear regression model and the extra sum-of-squares *F* test in GraphPad Prism. The majority of head and neck cancer cell lines do not form colonies from single cells, prohibiting clonogenic assays. A subclone of SCC-47 cells capable of forming colonies was isolated and used in all experiments. This subclone had similar levels of CIN to the parental cell line (Supp Fig. S7).

### Immunofluorescence microscopy

Immunofluorescence microscopy was performed as in (78). Briefly, cells were grown on coverslips and fixed in 4% formaldehyde, blocked, and stained with antibodies to α-tubulin (YL 1/2; 1:1000, Bio-Rad) and centromeres (HCT-0100, 1:1000, Cal Biotech), and counterstained with DAPI. Alexa Fluor-conjugated secondary antibodies (Invitrogen A21209 and A11013) were used at 1:200 for one hour at room temperature. Images were acquired on a Nikon Eclipse Ti2-E inverted fluorescence microscope using a Hamamatsu ORCA-FusionBT back-thinned camera or a Hamamatsu Orca Flash 4.0 camera and a 100×/1.4 numerical aperture (NA) oil objective. Images are maximum projections of 0.2-μm z-stacks deconvolved using Nikon Elements software.

### Murine studies

Animal studies were performed in compliance with relevant ethical regulations for animal testing and research. The study was approved by the Institutional Animal Care and Use Committee of the University of Wisconsin-Madison. 1 x 10^6^ FaDu cells in Matrigel were injected into each flank of 5 female athymic nude mice in each treatment group such that each mouse harbored two tumors. Tumors were allowed to grow until they reached approximately 200 mm^3^, at which time treatment was initiated. Mice were randomly assigned into 4 treatment groups. Unused docetaxel (20 mg/mL) was obtained from the UW hospital pharmacy per pharmacy procedures. Mice were injected with 2 mg/kg docetaxel I.V. (tail vein) and irradiation was performed 1 hour later. Mice were irradiated with 2 Gy daily for 5 days, mimicking patient treatment schedules. Animals were irradiated with a Precision Xray XRAD 320 with 1 Gy/minute delivered at 320 kV/12.5 mA at 50 cm FSD with a beam hardening filter with half-value layer of 4 mm Cu. The delivered dose rate was confirmed by ionization chamber. Mice were shielded with custom-built lead jigs to limit radiation exposure to the anterior 2/3 of the body. The sham irradiation cohort spent the same amount of time in the Xray treatment room as the irradiated group. One mouse (harboring 2 tumors) from each treatment group was sacrificed 24 hours after each treatment and portions of each tumor were immediately snap frozen or fixed in 10% formalin for 24 hours, dehydrated with 70% EtOH, then embedded in paraffin. 5 μm sections were cut, deparaffinized, and stained with anti-tubulin (DM1-α, Cell Signaling Technology #3873, 1:500) and anti-NuMA (NovusBio, NM500-174,1:100) antibodies overnight at 4°C. Samples were washed with PBS 3X and then stained with Alexa Fluor-conjugated secondary antibodies (Invitrogen at 1:200) and counter stained with DAPI (1:1000) for 2 minutes. The remainder of the mice were monitored and tumors were measured twice weekly until day 22 when they reached a size necessitating euthanasia. Tumor growth curves are shown as percent change in tumor volume (tumor volume at day 22/volume on day 4 of treatment*100) over time. Growth curves were compared using a mixed effect model. Head and neck cancer PDXs kindly provided by Dr. Randall Kimple were established as described (92) and treated with radiation or sham radiation as above. 5 μm FFPE sections were cut onto slides, and tissue was stained as above using anti-tubulin (DM1-α, 1:500), anti-centromere antibody (HCT-0100, 1:100, Cal Biotech), anti-pericentrin (Abcam, Ab4448, 1:200) and counterstained with DAPI. Tissue images were taken as described in the immunofluorescence microscopy section above.

### 6-centromere FISH and patient samples

Patients with a history of laryngeal cancer treated with definitive radiation therapy at University of Wisconsin were identified and biopsy samples were obtained under an approved minimal risk IRB protocol (2020-0011). 6-centromere FISH was performed at the Wisconsin State Laboratory of Hygiene which is a CLIA and CAP certified laboratory that routinely performs cytogenetic testing. Detection to enumerate chromosomes 3, 4, 7, 9, 10, and 17 was performed using FISH with the following 2 probe mixes in IntelliFISH hybridization buffer (Abbott Molecular, Des Plaines, IL): 1) Vysis CEP 3 (D3Z1) labeled SpectrumOrange (cat# 06J3613) localizing to 3p11.1-q11.1, Vysis CEP 7 (D7Z1) labeled SpectrumAqua (cat# 06J5427) localizing to 7p11.1-q11.1, and Vysis CEP 9 labeled SpectrumGreen (cat# 06J3719) localizing to 9p11-q11 in IntelliFISH hybridization buffer (cat# 08N8701); 2) Vysis CEP 4 labeled SpectrumGreen (cat# 06J3714) localizing to 4p11-q11, Vysis CEP 10 labeled SpectrumAqua (cat# 06J5420) localizing to 10p11.1-q11.1, and Vysis CEP 17 (D17Z1) labeled SpectrumOrange (cat# 06J3697) localizing to 17p11.1-q11.1. Localization of the probes was confirmed on pooled cytogenetically normal blood controls. Normal epithelial tonsil tissue (n=4 tonsils) obtained from the UW biobank served as a head and neck tissue control. After reviewing the corresponding H&E-stained tissue with a pathologist (R.H.), the hybridized area of interest was scanned using a BioView Duet Scanner (Rehovot, Israel). The number of copies of each chromosome was manually counted in at least 20 interphase nuclei for each probe set. Quantification of samples were performed in a blinded fashion. CIN was calculated as in (22). Briefly, the modal chromosome number for each chromosome was calculated and the average percentage of cells with non-modal chromosomes was averaged over the six chromosomes.

### Statistical Analysis

Statistical analysis was performed using PRISM version 10.2. Two-tailed Student *t*-tests were used to determine significant differences between groups unless otherwise indicated in the methods section or figure legend.

## Data Availability

The data generated in this study are available upon request from the corresponding author.

## Acknowledgements

We thank Heather Geye and Shari Piaskowski for their assistance with obtaining clinical samples, the University of Wisconsin Translational Research Initiatives in Pathology laboratory, supported by the UW Department of Pathology and Laboratory Medicine, P30CA014520, and NIH S10 OD023526 for its services, and the Weaver, Burkard, and Suzuki laboratories for insightful discussions. This project was supported in part by the Specialized Program of Research Excellence (SPORE) program, through the NIH National Institute for Dental and Craniofacial Research (NIDCR) and National Cancer Institute (NCI), grants P50DE026787 and P50CA278595 (SPORE PI P.M.H, DRP award to B.A.W.), R01CA234904 (B.A.W.), R01CA284747 (B.A.W.), R00CA160639, R37CA255330 (R.J.K), K08CA256166 (P.F.C), the Radiological Society of North America (Research Fellow Grant RF1904 to P.F.C), and the American Society of Clinical Oncology (Young Investigator Award to P.F.C).

## Author contributions

P.F.C. and B.A.W. designed the research; P.F.C., M.P., K.J., L.C.H., L. H., A.B., D.E., and E.B. performed experiments, analyzed data, and assisted with figure composition, E.M. assisted with microscopy figures, R.J.K., K.N. and J.W. provided reagents including PDX samples and data, P.M.H. assisted with the concept. P.F.C. and B.A.W. wrote the manuscript. All authors edited the manuscript.

