## Supplemental Figures for "Chromosomal instability increases radiation sensitivity"

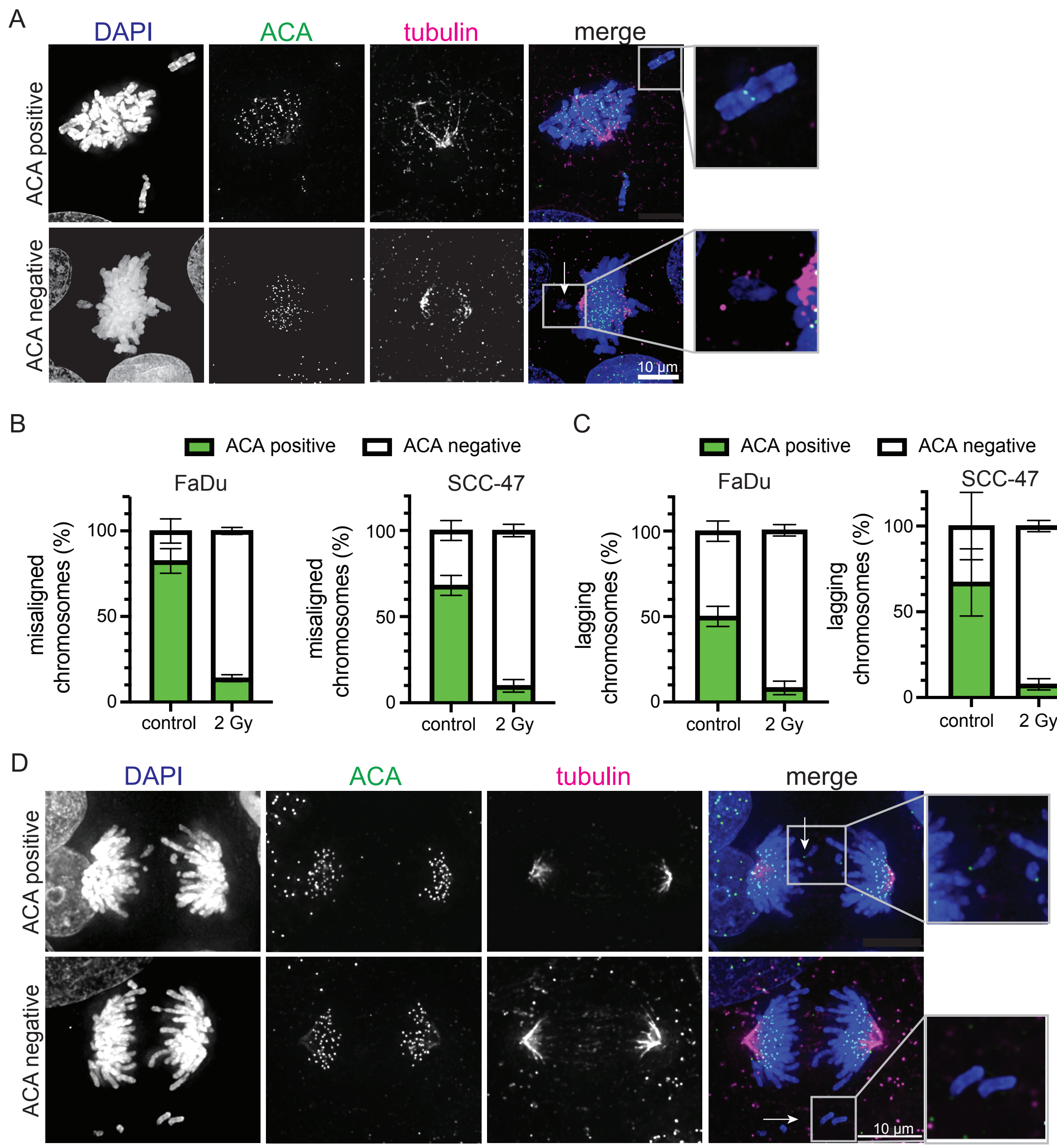


**Supplementary Figure S1. Radiation primarily induces structural CIN.** (A) Images of misaligned chromosomes with (top, inset) and without (bottom, inset) centromeres, identified with anti-centromere antibody (ACA). Loss of the centromere is evidence of structural chromosomal alterations. Arrow indicates misaligned chromosome without ACA staining. (B-C) Quantification of misaligned (B) and lagging (C) chromosomes with and without centromeres 24 hours after control treatment or 2 Gy of radiation. For the irradiated samples, n≥26 (range 26-113) misaligned and n≥40 (range 40-129) lagging chromosomes in each of 2 biological replicates. For control samples, n≥18 (range 18-31) misaligned and n≥15 (range 15-37) lagging chromosomes in each of 2 biological replicates. Error bars represent SD. (D) Images of lagging chromosomes with (top, arrow and inset) and without (bottom, arrow and inset) centromeres.

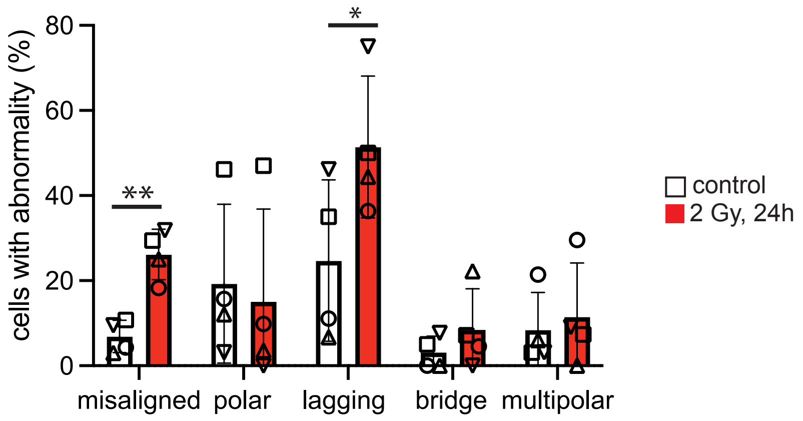


**Supplementary Figure S2. Radiation induces CIN in HPV-positive and HPV-negative head and neck cancer xenografts.** Quantification of mitotic defects in H&E-stained head and neck cancer xenografts at baseline (control) and 24 hours after 2 Gy of radiation. Each symbol represents a unique xenograft: triangle, SCC-2 (HPV-positive); inverted triangle, A253 (HPV-negative); square, SCC-47 (HPV-positive); circle, SCC-90 (HPV-positive). An average of 52 metaphases (range 22 – 71) and 15 cells in anaphase/telophase (range 8 – 22) were counted for each xenograft. * = $p<0.05$, ** = $p<0.01$ using 1-tailed t-test.

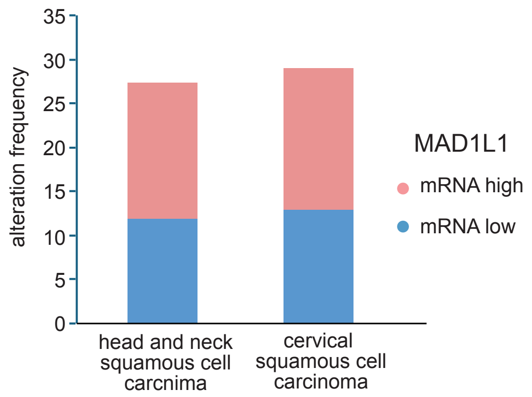


**Supplementary Figure S3.** ***MAD1L1*** **expression levels are commonly altered in head and neck and cervical cancers.** Expression of the gene encoding Mad1 (MAD1L1) is altered in 27% of head and neck squamous cell carcinomas (TCGA Firehose Legacy, 522 cases) and 29% of cervical squamous cell carcinomas (TCGA PanCancer Atlas, 248 cases). Data obtained from cBioPortal (93–95).

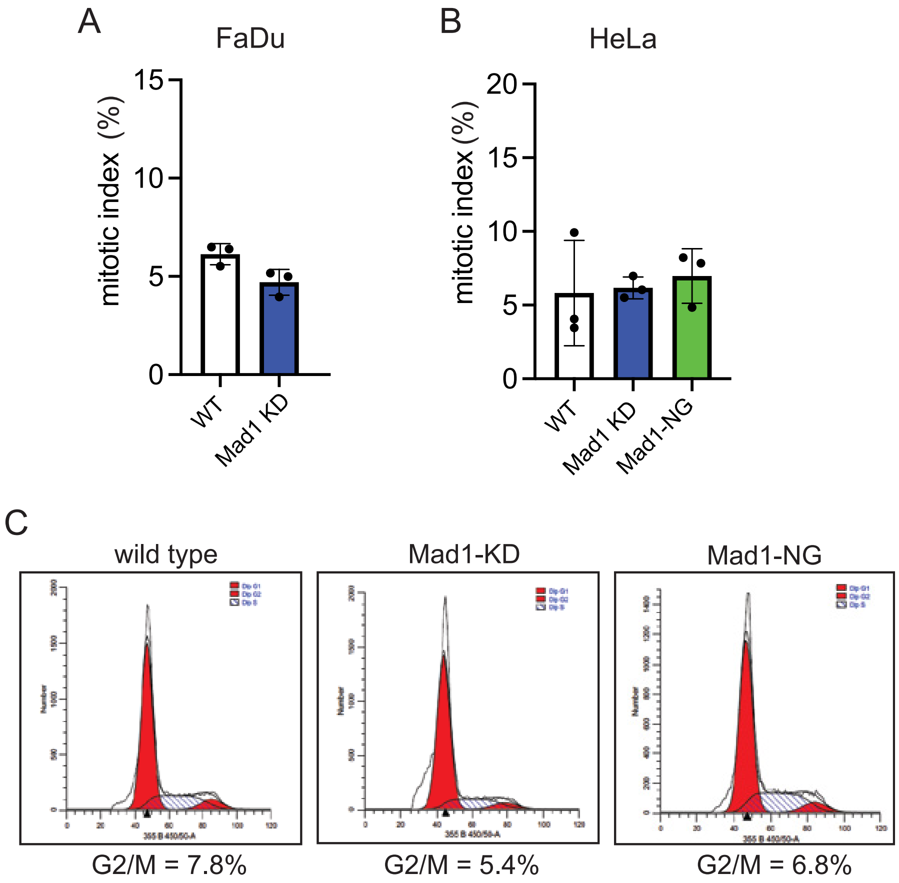


**Supplementary Figure S4. Altering Mad1 expression does not increase the fraction of cells in mitosis.** (A-B) Mitotic index (percent of cells in mitosis) remains unchanged in FaDu (A) and HeLa (B) cells after the indicated change in Mad1 expression. KD, knockdown. n=500 cells in each of 3 biological replicates. (C) Flow cytometry profiles showing that cell cycle distribution is unaffected by Mad1 knockdown or expression of Mad1-NG in HeLa cells.

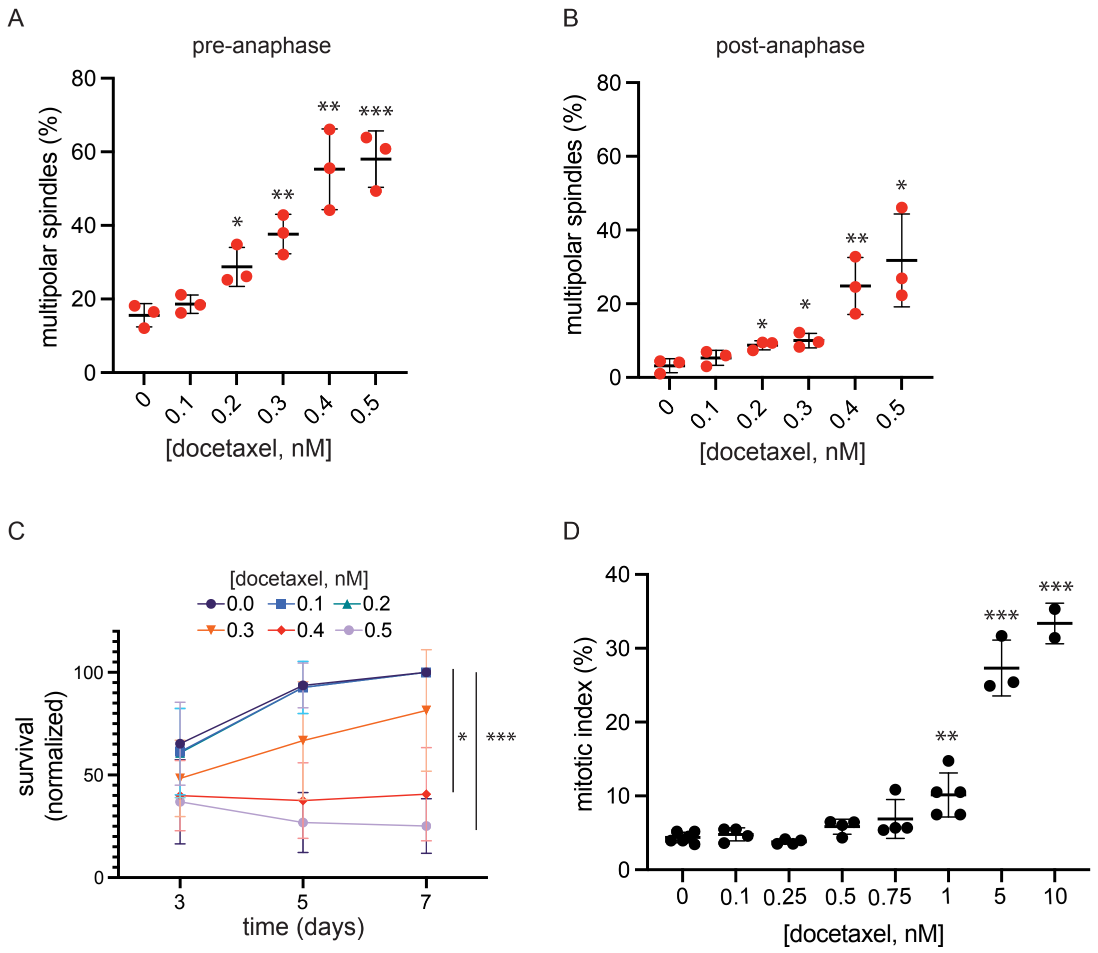


**Supplementary Figure S5. Docetaxel induces cytotoxicity without mitotic arrest by increasing chromosome missegregation on multipolar spindles that are maintained into late stages of mitosis.** (A-B) In HPV-positive SCC-47 cells, 24 hour docetaxel treatment induces formation of multipolar spindles in a dose dependent manner. Multipolar spindles occur in cells prior to anaphase onset (i.e. in prometaphase and metaphase, A) as well as in cells after anaphase onset (i.e. in anaphase and telophase; B). n≥100 cells in prometaphase+metaphase and 100 cells in anaphase+telophase in each of 3 biological replicates. (C) MTT assay of SCC-47 cells treated with increasing concentrations of docetaxel showing that concentrations that induce multipolar spindles in ≥20% of post-anaphase cells impair proliferation. n=3 biological replicates. (D) Concentrations of docetaxel that cause multipolar spindles and impaired proliferation do not cause mitotic arrest. Data obtained after 24 hours of docetaxel treatment. Error bars indicate SD. Statistical differences determined by 2-tailed t-test, * = p<0.05, ** = p<0.01, *** = p<0.001 versus DMSO.

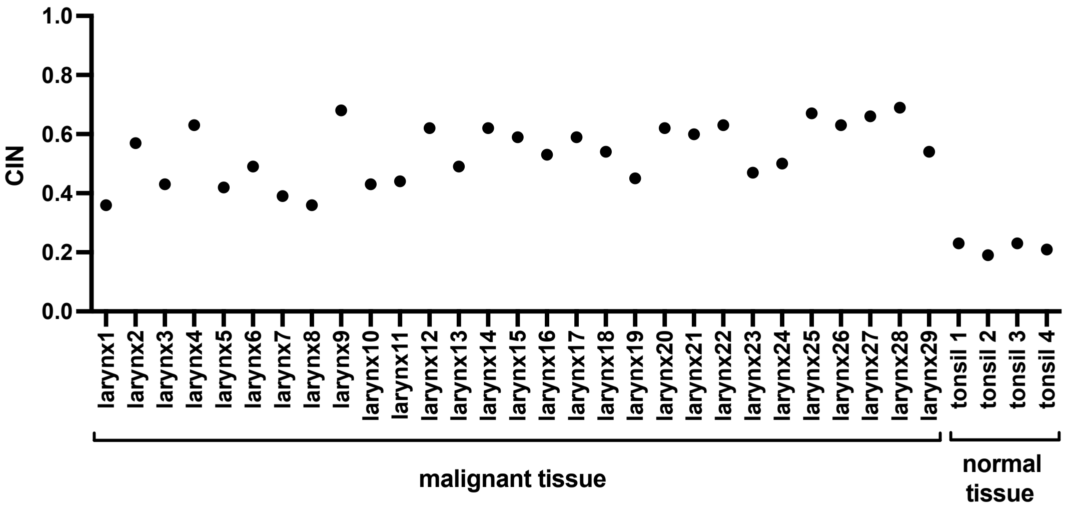


**Supplementary Figure S6. CIN in the laryngeal cancer patient cohort is heterogenous and higher than in control tissue.** CIN in each laryngeal tumor. Normal tonsil tissue was used as control. CIN was quantified based on the intercellular variability in copy number of 6 different chromosomes (see Methods). The observed rate of CIN in normal tissue is likely due to sectioning artifacts.

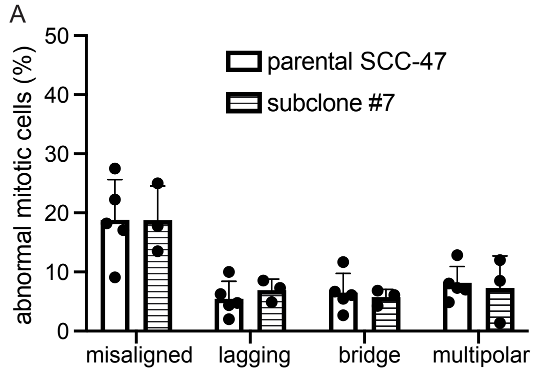


**Supplementary Figure S7. CIN does not differ between SCC-47 parental cells and the subclone used for clonogenic assays.** Quantification of mitotic defects that cause CIN in parental SCC-47 cells and the subclone capable of forming colonies. n≥100 cells in prometaphase+metaphase and 100 cells in anaphase+telophase in each of at least 3 biological replicates.
